# Spatially resolved distribution of pancreatic hormones proteoforms by MALDI-imaging mass spectrometry

**DOI:** 10.1101/2023.10.27.564470

**Authors:** Tháyna Sisnande, Felipe Lopes Brum, Daiane O. Matias, Fernando de Sá Ribeiro, Thayana Beninatto Moulin, Ronaldo Mohana-Borges, Mariana T. Q. de Magalhães, Luís Maurício T. R. Lima

## Abstract

Zinc plays crucial role in the immune system and endocrine processes. Dietary zinc restriction leads to the degeneration of the endocrine pancreas resulting in hormonal imbalance in β-cell. Proteostasis may vary depending on the stage of a pathophysiological process, motivating the development of tools aimed at the direct analysis of biological status. Among proteomics methods, MALDI-ToF-MS can serve as a rapid peptidomics tool from analysis of extracts or by histological imaging. Here we report the optimization of MALDI imaging mass spectrometry analysis of thin histological sections from mouse pancreas, allowing the identification of major islet peptide hormones and major accumulated precursors and/or proteolytic products of peptide hormones. Cross-validation of the identified peptide hormones was performed by LC-ESI-MS from pancreatic islet extracts. Mice fed a zinc-restricted diet had a relatively lower amount of peptide intermediates in comparison with the control group. These data provide evidence for complex modulation of proteostasis by an imbalance of micronutrients, directly accessed by MALDI-MSI.

## 1. INTRODUCTION

Diabetes Mellitus (DM) is a group of chronic diseases predisposing to other comorbidities and serious complications. DM can either be monogenic or, more commonly, polygenic, involving several environmental factors such as endocrine disruptors and macro- and micronutrient imbalances. It belongs to a group of endocrine and metabolic disorders characterized by deficient insulin and amylin production, secretion and/or response ^1^.

Diabetes mellitus type 1 (DM1) can be either autoimmune (DM1a) or idiopathic (DM1b). DM1a can be identified by autoantibodies to insulin (IAA), glutamic acid decarboxylase (GAD, GAD65), protein tyrosine phosphatase (IA2 and IA2β) and zinc carrier protein (ZnT8) ^2^. While multiple autoantibodies can predict future risk for DM1a (autoimmune type 1 diabetes) ^3,4^, high insulin levels in combination with high triglycerides anticipate insulin resistance and future risk for Diabetes mellitus type 2 (DM2) ^5^.

Dietary zinc restriction leads to the degeneration of β-cells associated with apoptosis and increased production of insulin and amylin^6^. This phenotype is consistent with a DM1b-like disease due to micronutrient imbalance, and, more broadly, motivates investigation of pancreatic proteostasis. However, insulin production may increase while pancreatic β-cell mass decreases, resulting in stable insulinemia ^6^. This lack of correlation between β-cell function and currently used plasmatic surrogates requires further investigation in the search for β-cell biomarkers.

Proteomics is a general definition for a wide range of powerful tools with applications in physiology, biomarker discovery, and targeted quantification. Proteomics can be applied to non-digested peptides or after proteolysis, which can be automated at the nanoscale using immobilized trypsin digest with peptide purification directly eluted in MS/MS analyzer ^27^, albeit with variations (up to 60%). As with any analytical approach, proteomics has several sources of uncertainty and limitations, such as large-scale identification of integer (whole) proteins and peptides, single *vs.* membranes, post-translational modifications, proteoforms, and more.

Several approaches can be used for targeted searches. Peptidomes can be obtained from complex biological matrices such as blood derivatives and tissues, using basic steps such as mechanical lysis and separation of the soluble fraction by centrifugation in the case of tissues (with or without solubilizing agents as detergents or urea/guanidinium), followed by solid-phase extraction with reversed phase (SPE-RP; C18 or C4 as preferred), elution and mass spectrometry measurements by Liquid-chromatography-Electrospray Ionization – Mass Spectrometry (LC-ESI-MS) or Matrix-Assisted Laser Desorption / Ionization – Time of Flight – Mass Spectrometry (MALDI-ToF-MS) are used. When the spatial distribution of biomolecules in tissues sections is required, MALDI-ToF-MS imaging (MALDI-MSI) of histological preparations can be performed ^7–11^.

Analytical proteomic approaches may have limitations, such as the use of denaturants, proteases, lack of quantitative robustness, and variability in extraction procedures (including liquid N_2_ freezing and manual mincing, bead homogenization, membrane extraction, and the protein precipitation). MALDI-MSI also has limitations, such as the inherent ionization properties of the molecules, matrix choice, sample preparation, time-consuming data collection, amino acid sequencing, and biased analysis. However, databases can help identify predefined desired molecules to support targeted proteomic analysis.

Robust analytical techniques, such as mass spectrometry imaging, has been applied to biological questions in recent years ^9,12^, including the pancreatic neoplasia model to identify potential new biomarkers ^13^ and the identification of insulin *in tissue* ^14^. MALDI-MSI has the advantage over single cells of preserving the spatial information and average heterogeneity of metabolic states of a single cell ^15^. However, optimization is still required for complex biological samples. Variables such as the choice of the matrix, the method of matrix application, the resolution, the type of tissue, the type of biological question, among others, are directly related to the quality of the results obtained.

Here we report the optimization of sensitivity of MALDI-MSI to map the peptidomic profile of pancreas islets, and its use in the identification of proteoforms ^16^ of the endocrine pancreas. This work provides advances in the MALDI-MSI as a powerful tool for peptidomic profiling in physiology or pathology and identify potential biomarkers for early endocrine dysfunction.

## 2. MATERIAL AND METHODS

### Material

EDTA-free Protease Inhibitor Cocktail (cOmplete™; Cat No. #11873580001) TEAB, 2-chloracetamide, α-cyano-4-hydroxycinnamic acid (CHCA), 2-mercaptobenzothiazole (MBT), Coomassie Brilliant Blue G250, collagenase type IV (CAS No. 9001-12-1) were obtained from Sigma-Aldrich Chemical Co. (St. Louis, MO, USA). Chromatography-grade solvents (acetonitrile, ACN; trifluoracetic acid, TFA; methanol; acetic acid) were obtained from Tedia Brazil.

### Ethics

This protocol was approved by the Institutional Bioethics Committee of Animal Care and Experimentation at the UFRJ (CEUA-CCS, UFRJ, #177/2019 and #045/2021).

### Peptidomic analysis by solid-phase extraction

#### Animals

Male Swiss mice (n=5) were weaned at 21 days of age onto standard chow^17^ (Nuvilab CR1; Nuvital Nutrientes S/A, Quimtia Brasil). After eight weeks, animals were euthanized by cardiac puncture, followed by blood and pancreas collection.

Peptidome mapping from pancreas and plasma was performed by extraction with macro and micro C-18 revered solid-phase extraction (SPE).

#### Pancreas extraction with macro-SPE

All procedures were performed on ice or at 4 °C unless otherwise indicated. The pancreas was minced with scissors and fragments of approximately 40 mg were transferred to polypropylene tubes (capacity 1.5 mL) containing 12.5 µL of lysis buffer (20 mM TEAB, protease inhibitor cocktail, and 40 mM DTT) per each milligram of tissue, and four zirconium oxide (ZrO) beads (2 mm each), and lysed with the aid of a bead mill (BulletBlender® tissue homogenizer; Next Advance™) for 5 min at room temperature with 5 min rest in ice and repeated. Samples were centrifuged at 15,000 x g at 4 °C for 30 min, 350 µL supernatant was diluted with 1.5 mL of 0.2 % TFA and subjected to extraction with Phenomenex® Strata® C18-E (55 µm, 70 Å, 200mg/3mL, Teflon® Tubes) according to product instructions, by loading the solutions into the column and collecting the flow-through in increments of 60 sec of centrifugation at 100 x g at room temperature into 15 mL polypropylene tubes.

Briefly, the column was activated with two steps of 2 mL 0.2% TFA/ACN, followed by two steps of 2 mL 0.2% TFA/water. The sample was loaded, washed with two steps of 2 mL 0.2% TFA/water, and the peptides were eluted with 1200 µL 70 % ACN/0.2 % TFA. The eluted material was aliquoted into 3 samples in low binding polypropypene tubes (nominally 1.5 mL) and dried in a vacuum concentrator (Vacufuge® plus; Eppendorf™) at room temperature without additional heating for about 3h. The dried samples were stored at −20 °C until resuspended with 100 µL water for further use. The yield was approximately 0.3 µg peptide/µL reconstituted solution (determined by Bradford assay).

MALDI-ToF-MS was performed with spots containing 2 µL peptide solution and 2 µL α-cyano (CHCA; 5 mg/mL in 50% ACN/0.3% TFA), and the 4 µL mixture was divided into 2 spots for technical duplicates.

#### MALDI-ToF-MS analysis

MALDI-ToF-MS was performed on an AutoFlex™ III spectrometer (Bruker Daltonics, Billerica, USA) calibrated with standards from the analytical facility (LMProt, CELAM, ICB, UFMG), in positive mode, using the method LP-ClinProt, in automatic mode, ranging from 1,500 to 20,000 m/z, suppressed at 1,400 m/z, with variable laser power up to accumulation of 7,000 shots 300 shots increment. Data were processed with FlexAnalysis™ v3.4 (Bruker, Germany).

### Matrix-assisted laser desorption/ionization - time-of-flight – imaging mass spectrometry (MALDI-ToF-MSI)

#### Animals

Swiss male mice were weaned at age 21 days onto either a control and a low-zinc diet as described ^6^. During the fourth week of the intervention, animals in the zinc cohort were sacrificed by cardiac puncture, followed by pancreatic sampling.

#### Sample preparation

For MALDI-MSI analysis, the pancreas was harvested and carefully placed in a plastic mold measuring approximately 1.0 cm x 0.5 cm x 0.5 cm. All organs were oriented in the mold so that the pancreatic duct was visible, and the posterior islet could be optimally located. The organs were individually covered with 10% warm gelatin solution (Cat # 9000-70-8, Sigma-Aldrich, USA) in ultrapure water, at 37 °C. Immediately, the molds were placed in a bath of solid carbon dioxide (dry ice) with acetone until completely frozen, transferred to falcon tubes, and again bathed in a solid carbon dioxide/acetone bath. The samples were stored at −80° C until the sectioning procedure. The organs have thus been acclimated at −20 ° C for at least 4 h, and sections were subsequently performed. Sections of 10 µm at −20 ° C were performed in cryostat (Leica CM1860UV, Leica Biosystems, Germany). The first section was stained with toluidine blue for visual inspection of the presence of islets. The next section was allocated to slides containing indium tin oxide (Cat No. #8237001; ITO - Bruker, USA) suitable for the MALDI-MSI technique. The tissues sections were collected and analyzed by MALDI-MSI followed by staining with toluidine blue solution to overlay the images facilitating the localization of the islets. Few freshly made staining solutions (1% toluidine blue and 2% borate in water) were dropped on the slide, warmed on a hot plate, washed with water, and photographed. The sample slices were washed 3 times with a solution of methanol:chloroform (1:1) for 30 s each time and air dried between each wash. The slides were then completely dried in a desiccator for approximately 15 minutes.

#### Matrix dispersion

Sample analyses were performed using either 4-hydroxycinnamic α-cyano acid (CHCA - Cat No. #201611-92-9, Sigma-Aldrich, USA) or 2-mercaptobenzothiazole (MBT – Cat No. #149-30-4, Sigma-Aldrich, USA) as the matrix in two different protocols.

First, CHCA (100 mg) was dissolved in acetone (q.s.ad.), dripped on the sample, allowed to dry in a desiccator for about 15 min, and the quality of signal and imaging of pancreatic peptides was evaluated.

For the second protocol, MBT (300 mg) was dissolved in acetone (q.s.ad.) followed by solvent evaporation on the sublimation apparatus for matrix crystallization apparatus as described elsewhere ^19,20^. Afterwards, inside the sublimation, the sample remained for 3 min at 50 mTor, at 180 ° C for deposition of the matrix by sublimation.

#### Data collection

Tissue sections were subjected to data collection using MALDI-ToF-MS (Autoflex Speed maX, Bruker, USA, belonging to CEMBIO-UFRJ), in positive mode, ranging from 500 to 6000 m/z, with medium focus and laser power at 50%, 200 shots/image collected with a lateral resolution of 50 µm.

#### Data processing

The generated images were analyzed using FlexImaging® (version 4.1; Bruker, Germany; licensed to CEMBIO-UFRJ). Processing included smoothing (Savitzky-Golay; width 5 m/z, 5 cycles) and baseline subtraction (TopHat).

#### MALDI-FT-ICR-MSI

The FT-ICR Solarix XR 7T spectrometer (Bruker Daltonics) collected the high-resolution MALDI-FT-ICR-MSI. The slides prepared as previously described (control animal; internal code LSC1A3) with MBT matrix were subjected to MALDI in positive mode, in the range of 500 to 6000 m/z, with laser saturation at 45%, with 500 shots/frame, time domain data set size of 1 MW, and resolution of 29,000 for 3,121 m/z. Data was converted using msConvert from ProteoWizard ^21^, and peptide fragments were prospected with PepSeq (MassLynx, Waters) and mMass® (http://www.mmass.org/) ^22,23^. The peptides were annotated based on their monoisotopic mass as calculated using PeptideMass (https://web.expasy.org/peptide_mass/) with their corresponding sequences obtained from UniProt.

### Peptide identification in the islets

#### Islet isolation

The procedure was conducted with the pancreas collected from the control mouse described above receiving the standard AIN-93 diet. All procedure was performed using Hank’s buffer (0.14 M NaCl, 5 mM KCl, 1 mM CaCl_2_, 0.4mM MgSO_4_-7H_2_O, 0.5 mM MgCl_2_-6H_2_O, 0.3 mM Na_2_HPO_4_-2H_2_O, 0.4 mM KH_2_PO_4_, 4 mM NaHCO_3_). Isolation was performed with modifications as described elsewhere ^24^.

The liver duct of euthanized mouse was blocked with surgical line and with the hemostatic scissors blocked the duct that goes to the intestine. Carefully, with the duct well stretched, the pancreatic duct was cannulated and perfused with collagenase type IV at 1.0 mg/mL. The inflated pancreas was removed, minced in glass petri dish with Hank’s buffer, transferred to a polypropylene tube with 1 mL collagenase type IV (1 mg/mL Hank’s buffer), and incubated at 38° C for 30 min. The digestion was transferred to ice bath, added equal volume of cold Hank’s buffer and the islets were pelleted by centrifugation at 4° C. The supernatant was removed and the islets were Hand-picked with aid of a pipet, with the following methodology_25,26._

#### Liquid-chromatography mass spectrometry (LC-MS) analysis

Isolated islets were dispersed in Hank’s buffer containing 3 mM PMSF and sonicated (5 min, 75% A, pulse 15 s, interval 10 s; Vibracell VCX130, Sonics & Materials, CT, USA). Posteriorly, the lysate was filtrated in Centricon™ 10 kDa (Merck Millipore). The flow-through was loaded into SPE-C18 (MacroSpin Columns, Harvard Apparatus) previously activated with 400 μL TFA 0.1% in 50% ACN followed by 800 μL 0.1% TFA in water. The resin was washed 5 x 400 μL 0.1% TFA in water and peptides were eluted with 400 μL 50% ACN/0.1% formic acid followed by 400 μL 90% ACN/0.1% formic acid. The fractions were combined and dried out with vacuum concentrator at room temperature. The sample was reconstituted with 40 μL 0.1% formic acid with 3% ACN and subjected to analysis in a UPLC Nexera X2 (Shimadzu) with Ascentis column C18 (150 x 1 mm, 3 μm particle size, Supelco), with oven at 50 °C, flow 65 μL/min, injection volume 10 μL, with the following gradient (A = 0.1% formic acid in water, B = 0.1% formic acid in ACN): 0-1 min, B=2%; 3 min, B=8%; 40 min, B=25 %; 55 min, B=50%; 58 min, B=50%; 59 min, B=95%; 71 min, B=95%; 72 min, B=2 min; 85 min, B=2%.

Detection was performed with Maxis Impact, ESI-Q-TOF (Bruker Daltonics), using sodium formiate (100 μM in water:isopropanol 1:1) as calibrant. The acquisition was performed in DDA/AutoMS, with isolation/fragmentation of 3 precursors / cycle, in a scan range of 50 - 1500 m/z, 1 Hz acquisition rate, with positive ESI mode, nebulizer set at 1.2 bar, dry gas at 8 L/min and dry Temperature at 200 °C. The capillary voltage was set at 4500 V, and the end plate offset at −500 V with quadrupole low mass set at 300 m/z. Data were analyzed with Compass DataAnalysis (Bruker, Germany).

## 3. RESULTS

### Peptidomic of whole pancreas by C18-SPE / MALDI-ToF-MS

The whole pancreas peptidome of adult mice on a standard control diet (AIN-93) was performed. After euthanasia, fragments of pancreas were lysed in the presence of protease inhibitors, the supernatant was fractionated in reversed-phase (C18) solid-phase resin (SPE), and the resulting extract was subjected to MALDI-ToF-MS evaluation. A complex spectrum was obtained with ions with varying intensity. The intact C-peptide of insulin 1 (INS-1), the chain B of both INS-1 and INS-2 (**Fig. 1**) were identified by directly examination of the ion list. Glucagon, amylin, polypeptide P (PP), somatostatin-28, C-peptide of INS-2 and the chain A of insulin could not be identified (**Fig. 1**). Direct extraction from the pancreas and offline SPE-C18 peptide partitioning did not allow the identification of the major peptide hormones, indicating a limitation of this approach in profiling the endocrine pancreas.

**Figure 1.**
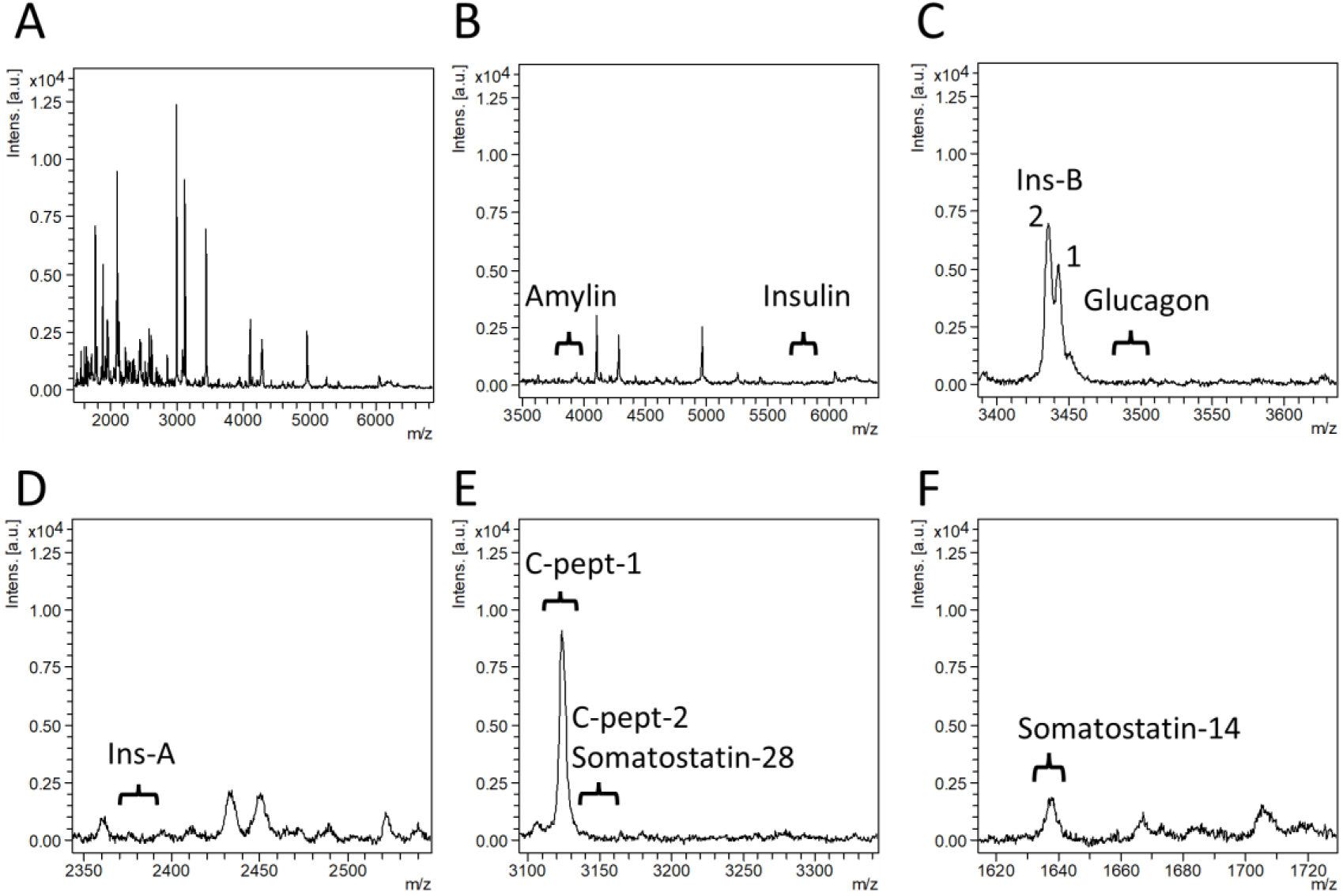
Peptidomic of mouse pancreas extract by SPE-C18 / MALDI-ToF-MS. Data show the different ranges and peaks observed in the mass spectra of mouse pancreas (n=5) extracts obtained using reversed phase (SPE-C18, 70 % ACN fraction).

A) full spectra
B) insulin and amylin range;
C) Insulin chain B and glucagon range;
D) Insulin chain A
E) C-peptide and somatostatin-28 range;
F) somatostatin-14.

### Pancreas MALDI-MSI standardization

The first attempts to perform MALDI-MSI with pancreas were conducted using CHCA as the matrix. This approach allowed the identification of a few ions (**Fig. 2**). The highest peak plotted image (Fig. 2A) shows the location of the islets, our principal region of interest (ROI 1), and the lack of the signal represents a region outside the islet (ROI 2). The ROI 1 region plotted spectra shows the highest peak, corresponding to the annotated insulin, with a near main peak and without further clear peaks in the range of 1,000 to 8,000 m/z (**Fig. 2B**). The spectra from the ROI2 without the corresponding annotated insulin peak, were mainly dominated by noise (**Fig. 2C**). To optimize *in-situ* extraction and measurements of other peptides, the same slice was subjected to matrix recrystallization with water:ACN (1:1) in the presence of acetic acid (5% v/v). The mass spectrum resulting from the recrystallized sample showed greater ion complexity compared with the first assay (**Fig. 2D**), allowing the identification of ions with m/z compatible with [M+H]^+^ of insulin (5817.3m/z), amylin (3920 m/z), INS-1 chain A (2366.98 m/z), INS-1/2 chain B (3439.2 m/z), INS-1 C-peptide (3123.0 m/z) and other high-intensity non-identified ions (2994.0 m/z, 4974.0 m/z and 6140.2 m/z). Despite the improvements obtained with recrystallization, CHCA was not a good choice of the matrix because only a small number of ions were detected, and lacked signals compatible with other expected peptides from the endocrine pancreas.

**Figure 2.**
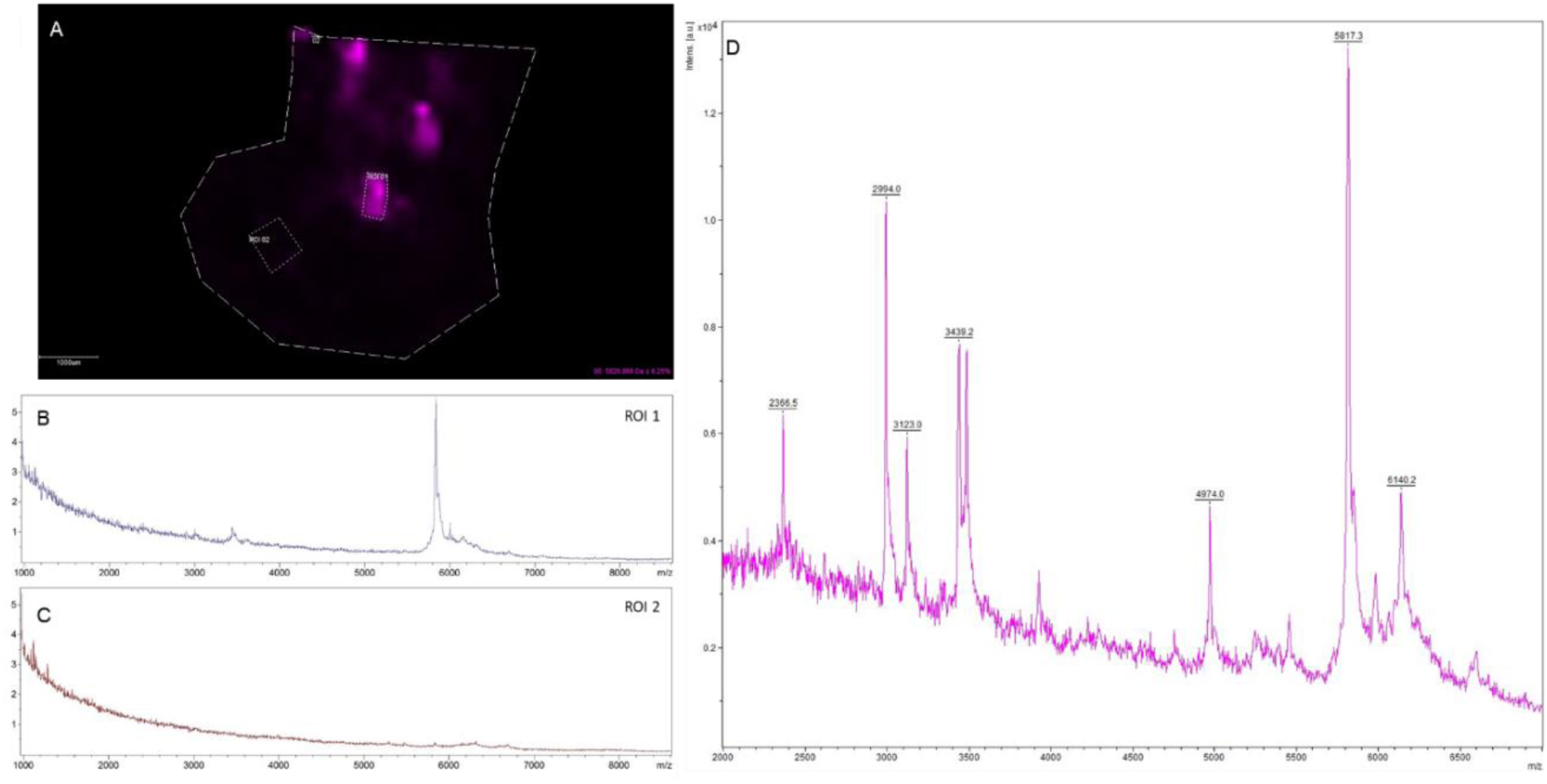
Analysis of α-cyano 4-hydroxycinnamic acid as a matrix for MALDI-MSI of pancreas. Pancreas was dripped with a-cyano 4-hydroxycinnamic acid solution in acetone. In A, 2D mapping for the highest intensity ion (5821 kDa - purple). In dotted lines are delimitated two regions of interest (ROI). B) mass spectrum of ROI-1. C) mass spectrum of ROI-2. The previous slide (A) was subjected to recrystallization with a solution of H_2_O: ACN (1: 1) and reanalyzed, and the mass spectrum of ROI-1 is shown in (D).

An alternative sublimation method used 2-mercaptobenzothiazole (MBT) as matrix. A large number of the major known peptides from the endocrine pancreas were detected by sublimation of MBT using the matrix coating protocol (**Fig. 3**). Several tissue regions compatible with pancreatic peptides as shown by staining with toluidine blue (**Fig. 3A**), yielded similar regions of interest (ROI) (**Fig. 3B**) with a high-intensity mass spectrum (**Fig. 3C**). These ROI showed ions compatible with pancreatic peptides as [M+H]^+^ and Na^+^ or K^+^ adducts (**Fig. 3D**) including insulin, C-peptide, amylin and glucagon, as identified based on their sequence and average mass as annotated in UniProt (**Table 1**). A clear distinction between INS-1 and INS-2, or between C-peptide-1 and C-peptide-2), was not possible based on spectral resolution. Instead, the different islets showed abundance, specificity and consistent mass spectra that were considered satisfactory and therefore used for further MALDI-MSI measurements.

**Figure 3.**
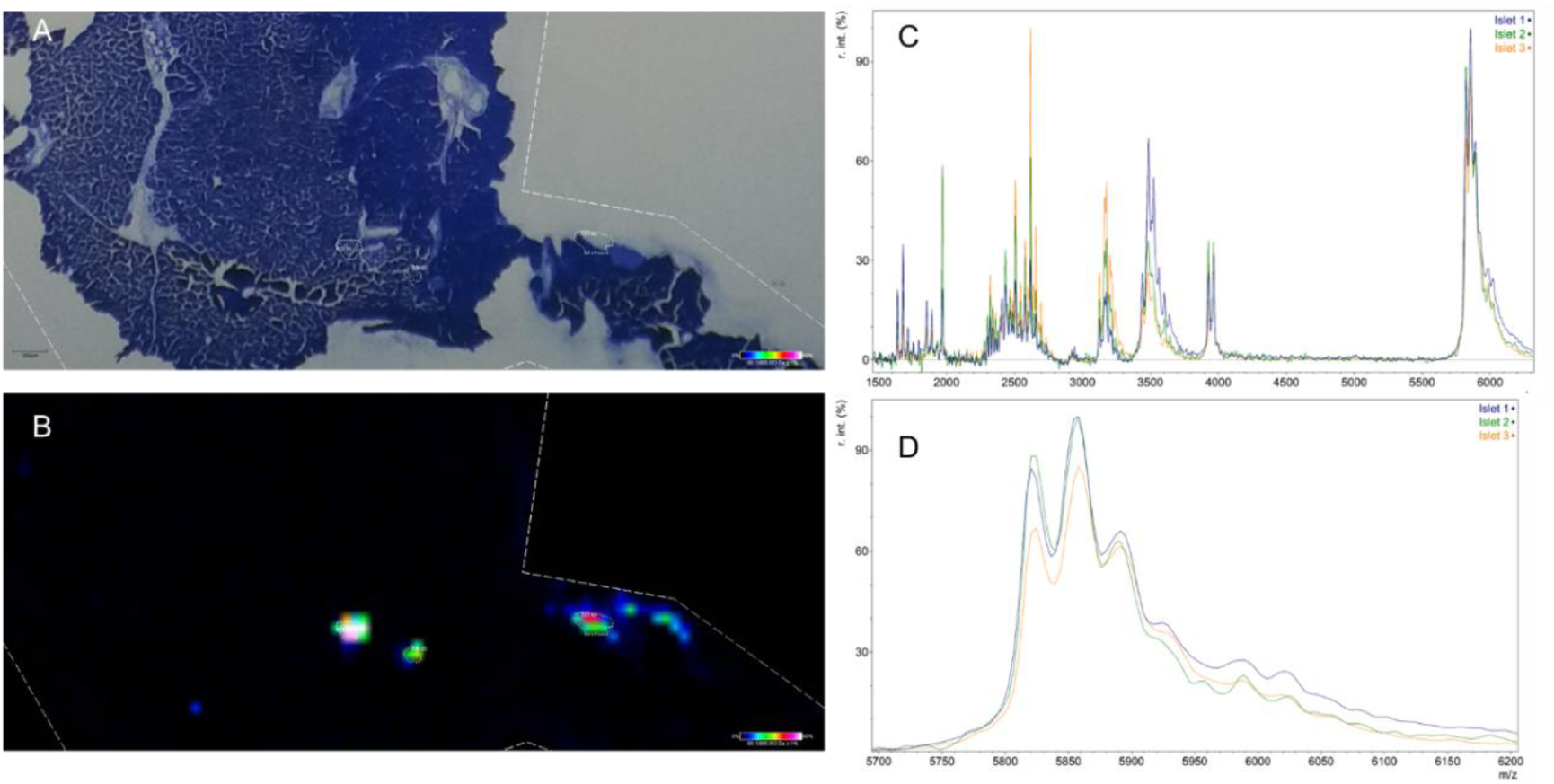
Analysis of 2-mercaptobenzothiazole as a matrix for MALDI-MSI of pancreas. **(A)** tissue from control animal stained with toluidine blue; **(B)** assignment of ROI for 5866 m/z (insulin as potassium adduct, as shown in inset, panel **D)**; **(C)** mass spectrum of three regions of interest selected as shown in **(A)** (colors correspond to different islets); **(D)** expanded view of **(C)** for the insulin region, evidencing three main peaks possibly compatible with free INS-1, and K and Zn adducts.

**Table 1.**
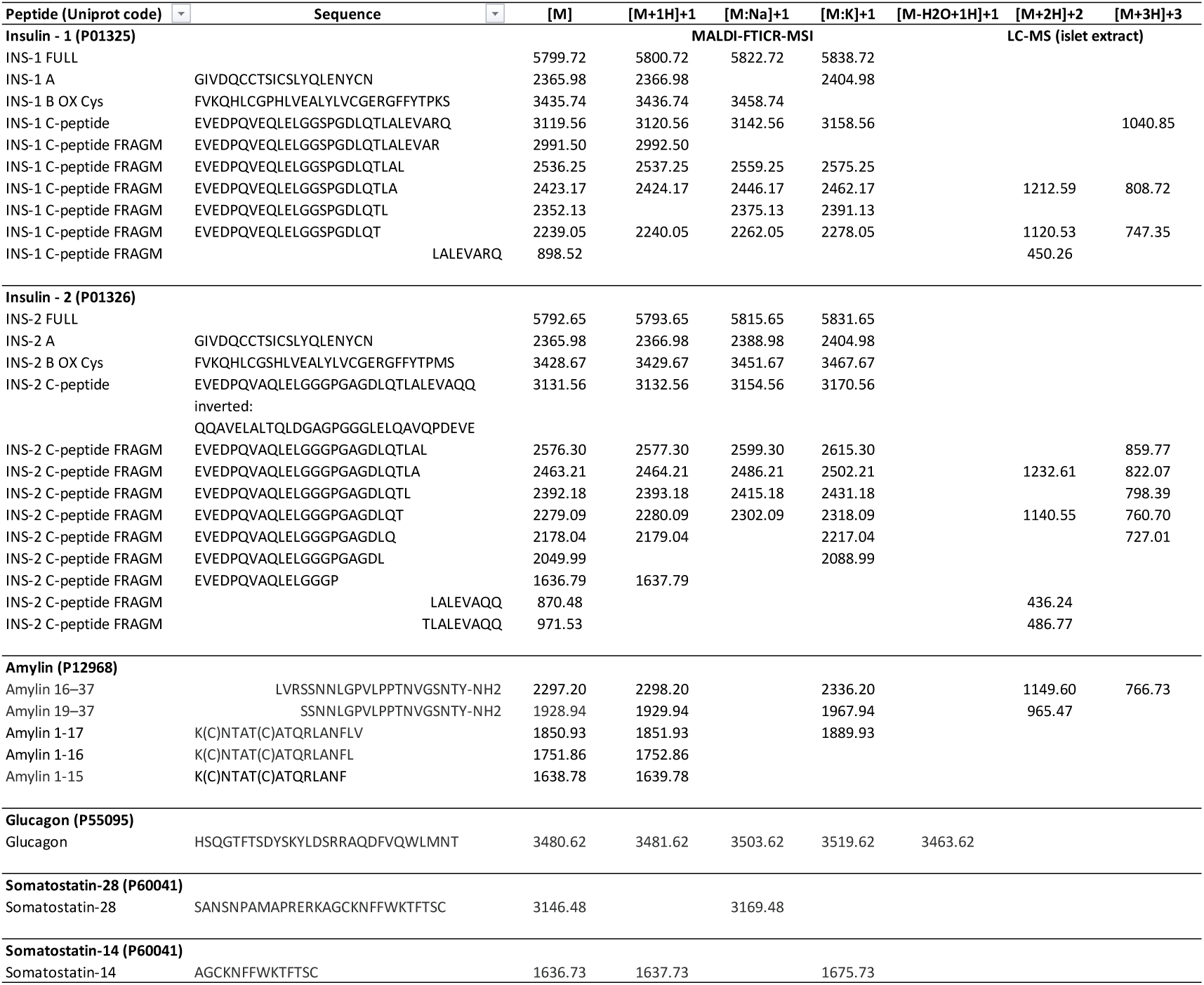
Ions identified in MALDI-MSI-FTICR from islets in histologic slices, and in undigested LC-MS-MS from islet peptide extracts of mouse pancreas. List of pancreatic peptides identified MALDI-FT-ICR (native or Na/K adduct) of pancreatic islets, as deduced from their monoisotopic mass calculated using PeptideMass (Expasy) using amino acid sequence listed in UNIPROT

### Differential mapping of pancreatic hormones using MALDI-MSI

Differential peptidome mapping directly from histological sections of pancreas was performed using the MBT-matrix protocol, as it was found to be more suitable and reproducible. For this reason, we proceeded with all sample preparation using this protocol. We measured the matrix deposited per area as at about 0.2 μg/cm^2^. Improvements were observed by recrystallizing the dispersed matrix in steam/atmosphere using water:ACN at a 1: 1 ratio, plus 5% acetic acid, for 1 minute at 60 ° C.. Measurements were performed with constant parameters to allow direct intensity comparison. The MALDI-MSI distribution profile of insulin (**Fig. 4A**), C-peptide (**Fig. 4B**), amylin (**Fig. 4C**), glucagon (**Fig. 4D**), and somatostatin (**Fig. 4E**) were based on the ion with the highest intensity for the corresponding adduct. Ions corresponding to these peptides were found colocalized, demonstrating the specificity of the method. Insulin was detected in well-defined regions (**Fig. 4A**). At the same time, C-peptide (**Fig. 4B**) and amylin (**Fig. 4C**) were also found in some regions adjacent to the ROI for the low-zinc group, which could be due to lixiviation during sample preparation and/or other unknown reasons. While insulin, C-peptide, amylin and glucagon showed high intensity in islets in both groups, the signal from somatostatin was low in the ROI under analysis (**Fig. 3E**), which could be due to either the amount of this hormone or methodological problems.

**Figure 4.**
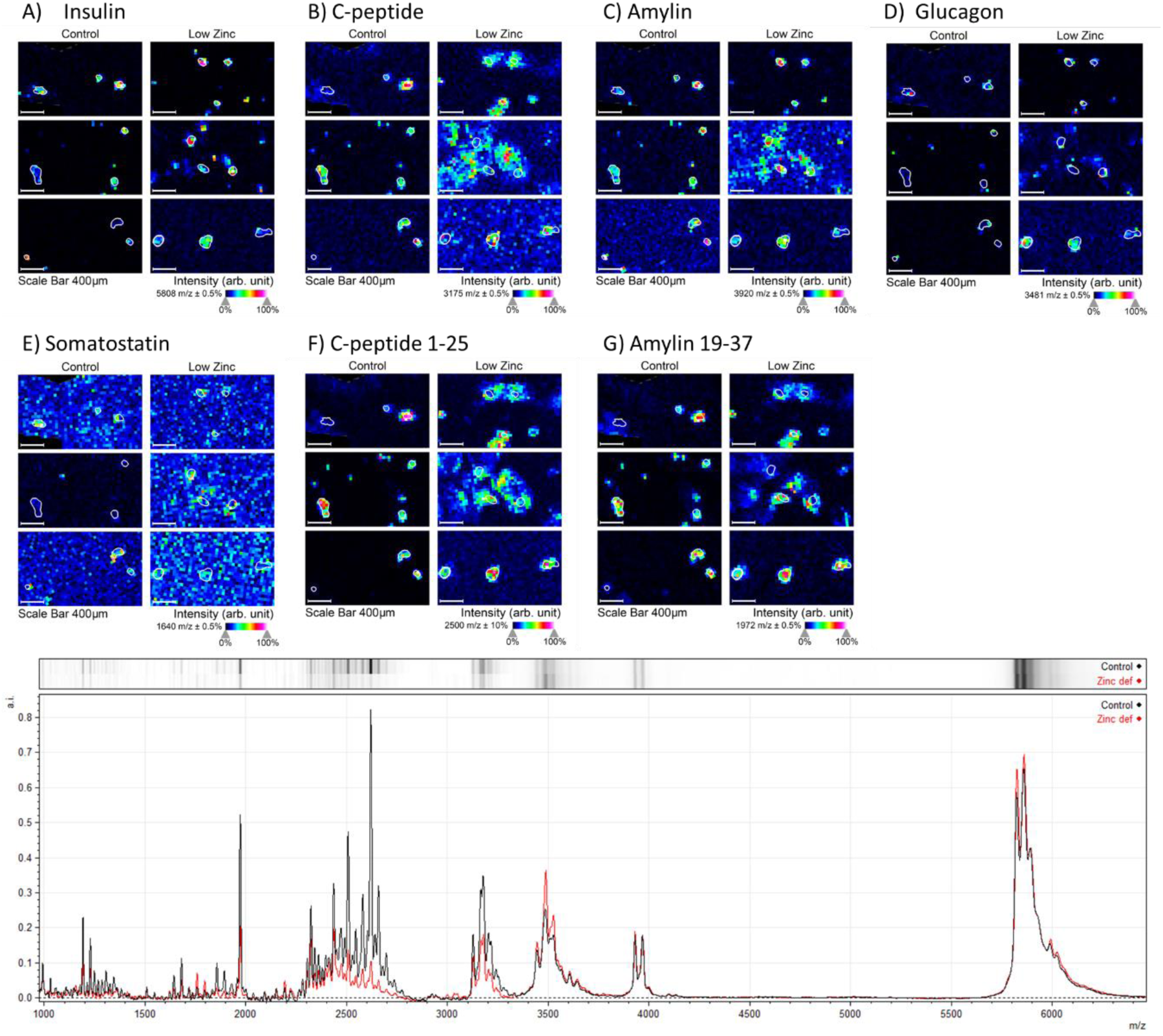
MALDI-IMS in control and zinc-restricted groups. Images were generated for **(A)** insulin 5808 m/z ion, **(B)** C-peptide 3175 m/z adduct, **(C)** amylin 3920 m/z ion, **(D)** glucagon 3481 m/z ion, **(E)** somatostatin 1640 m/z ion, **(F)** 2500 m/z ion (K-adduct of C-peptide from INS-2, fragment aa 1-25) and **(G)** 1972 m/z ion (K-adduct of amylin fragment aa 19-37). The color scale of the images was plotted based on the intensity of the signal (from black to white) generated by the mass spectra for each ion detected, with black being the unsigned spectrum for each m/z and white being the highest intensity signal for the same m/z relationship. N=3/group. **(G)** Comparative analysis of the mass spectra of the control and intervention groups. Three islets - ROIs - of each animal, from three animals (total 9 ROI/group) were analyzed, generating 1 spectrum. This spectrum was considered for each animal (N = 3 / group) and the average for the group was performed. In black, average of the 9 spectra in the control group. In blue, average of 9 spectra of ROIs in the zinc-restricted group. Highlighted peaks m/z 5808 - insulin; m/z 3920 - amylin; m/z 3482 - glucagon; m/z 3176 - C-peptide; family m/z ∼2500; m/z 1972 and m/z 1641 - somatostatin.

The average spectra of the nine ROI for each group (**Fig. 4**) show the similar intensity of insulin, amylin, and glucagon ions (native and adducts). In contrast, C-peptide-related ions of approximately 1972 m/z and a group in the range of 2500 m/z were lower in the zinc-restricted group.

Individual ROI of three animals from the control group and three animals from the group with zinc-restriction show similar ion distribution profiles with high intensity, although with differences between ROI in the same animal or even in the same group (control *vs.* intervention; **Fig. 5**). These results suggest independent behavior of individual islet, highlighting the importance of methods such as mass spectrometry imaging in characterizing biological individualities from a proteostasis point of view.

**Figure 5.**
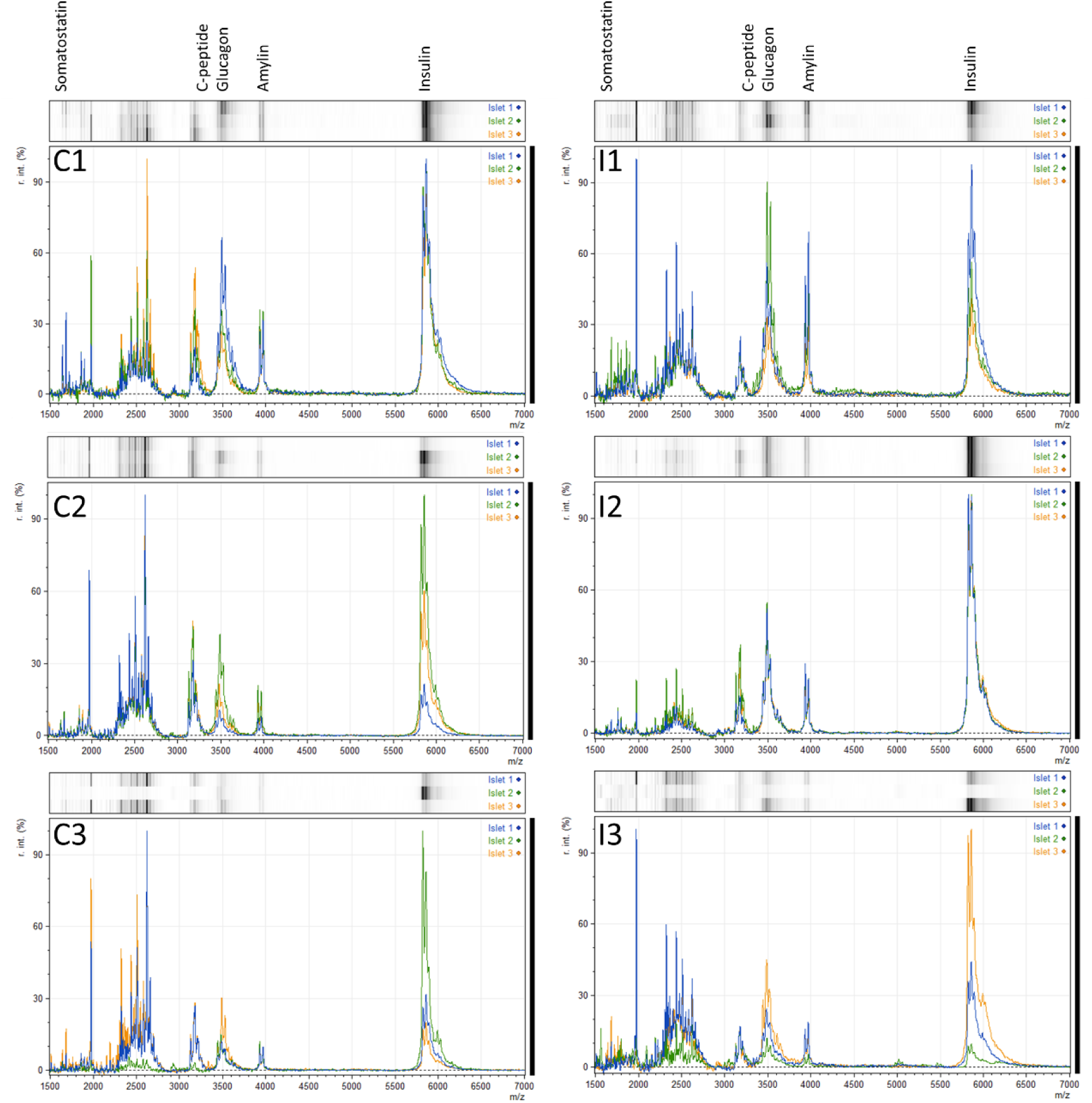
Homogeneity in peptide distribution among islets. Spectra from three islets (different colors) from each animal were obtained and compared among three animals/groups: control (C1, C2 and C3) and dietary zinc-restriction intervention (I1, I2, I3).

### Identification of proteoforms of hormone peptides

All spectra from islets revealed the prevalence of other unidentified high-intensity ions, including a strong ion at about 1972 m/z and a group of intense ions near 2500 m/z (2,200 – 2,700 m/z; **Fig. 3**, **Fig. 4**, **Fig. 5**). These ions were found in all spectra of each ROI (**Fig. 5**) and were colocalized in the other peptide hormones shown here in the same ROI (**Fig. 4**).

To gain insight into these ions’ nature and confirm the identity of the known major hormone peptides previously identified, a high-resolution spectrum was obtained from an islet by MALDI-FTICR (**Fig. 6**; **Table 1**). The monoisotopic mass of the major peptide hormones of the islet was confirmed, INS-1 and INS-2 (**Fig. 6A**), C-peptide-1 and C-peptide-2 (**Fig. 6B**), amylin (**Fig. 6C**), glucagon (**Fig. 6D**), somatostatin-28 (**Fig. 6E**) and somatostatin-14 clearly identified (**Fig. 6F**). The ion with an average mass charge of 1972 m/z indeed proved to be a potassium adduct, corresponding to [M+K]^+^ with exactly 1967.93 m/z, starting from the original [M+H]^+^ with 1929.97 m/z. The amylin fragment [19–37]amylin proved to be compatible with the identified ion, whereas the complementary fragment [1–18]amylin (containing an ^18^Arg) could not be detected. Other amylin proteoforms were also identified in the MALDI-FTICR spectra, such as [16–37]amylin, [1–17]amylin, [1–16]amylin, and [1–15]amylin (**Fig. 7**; **Table 1**).

**Figure 6.**
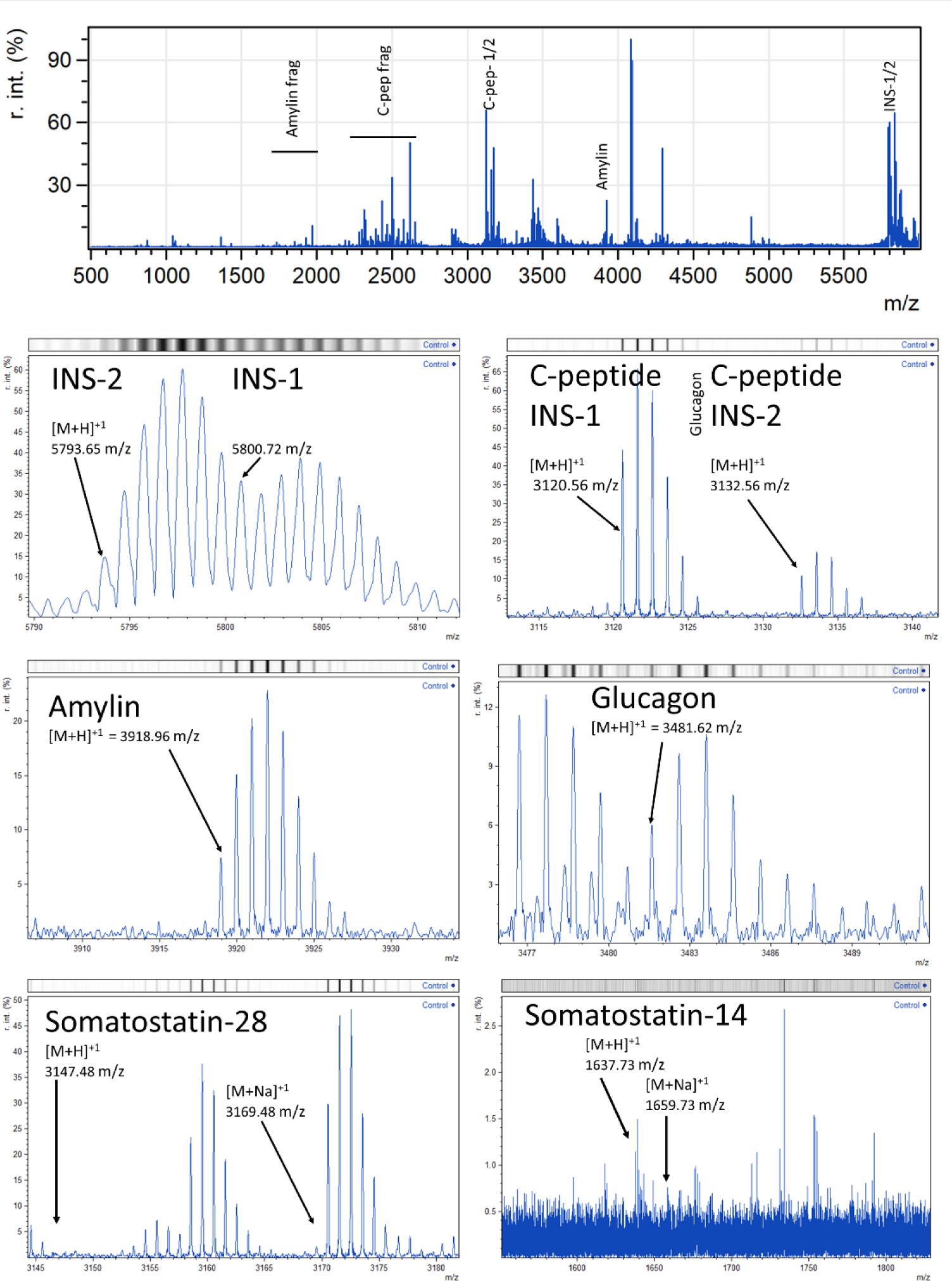
High resolution MALDI-FTICR. Characterization of the peptides present in the regions of interest of the control animal 1. Individual mass spectra of each region of interest selected in the original image, in the abscissa mass/charge ratio and in the ordered arbitrary intensity. Blue, islet 1. Green, islet 2. Orange, islet 3. Highlighted peaks m/z 5808 - insulin; m/z 3920 - amylin; m/z 3482 - glucagon; m/z 3176 - C-peptide; family m/z 2500; m/z 1972 and m/z 1641 - somatostatin.

**Figure 7.**
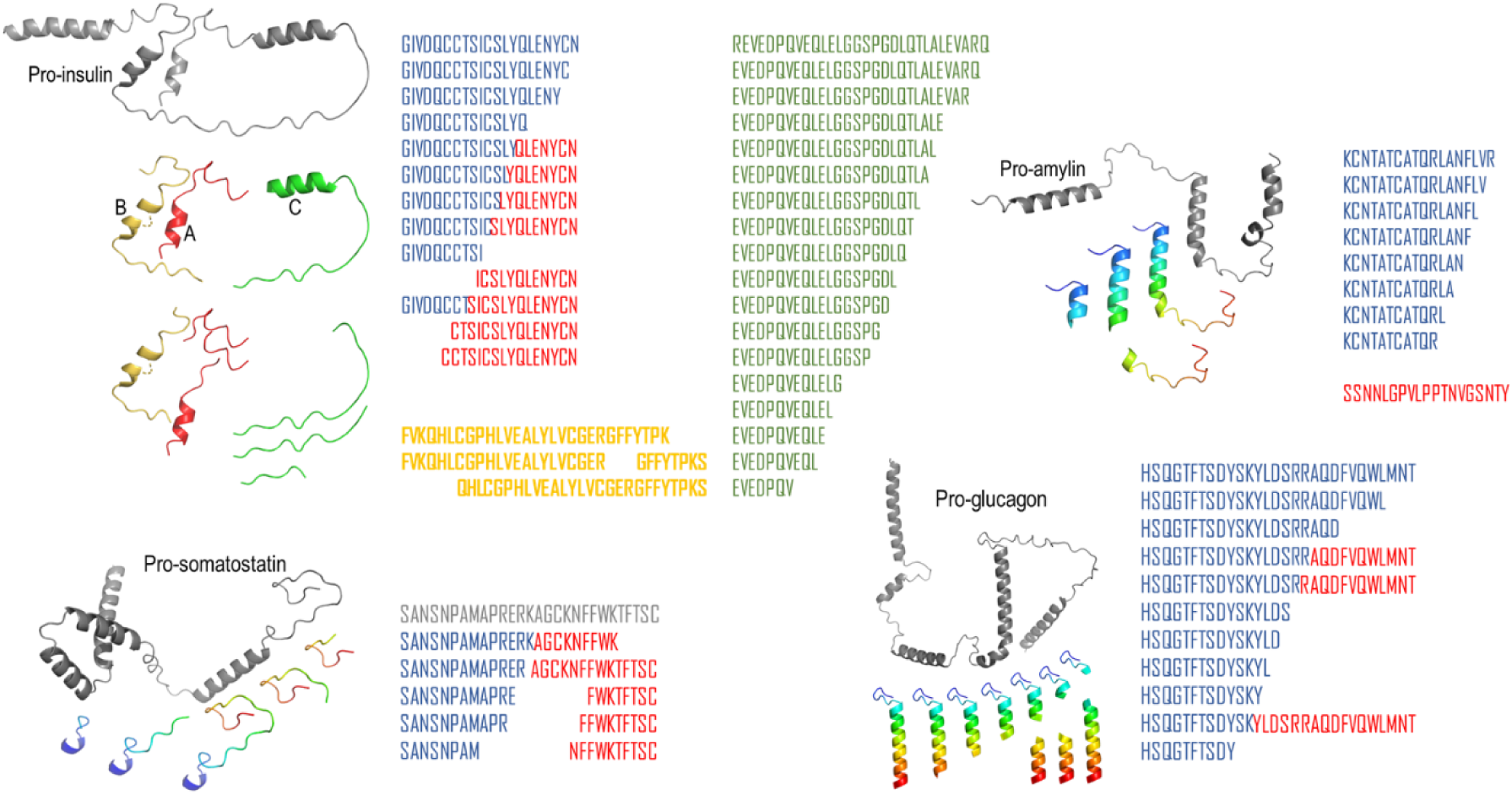
Schematic representation of progressive degradation of main pancreatic hormones. The sequences correspond to selected fragments (different colors) of the mature hormone peptides from mouse reported in the *ProteomeXchange* Consortium (https://massive.ucsd.edu/). The predicted structures of the pro-hormones (grey) were obtained from *AlphaFold* at UNIPROT (access 05/05/2023) for insulin-1 (P01325), amylin (P12968), glucagon (P55095) and somatostatin (P60041). The colored structures are representations of some fragments identified.

The m/z family group of 2500 was found to include the proteoforms of C-peptides (**Fig. 7A**; **Table 1**) as [M+H]^+^ and adducts [M+Na]^+^ and [M+K]^+^. Ions corresponding to proteoforms of the C-peptides were identified for [1–21]C-peptide-1 to [1–24]C-peptide-1 in the variation step of one amino acid and for [1–22]C-peptide-2 to [1–26]C-peptide-2 also in variation step of one amino acid (**Table 1**). Ions corresponding to other fragments of C-peptide-1 or C-peptide-2 with decreasing size toward the N-terminus and the expected complementary peptides for the total C-peptides were not found in the MALDI-FTICR.

### Confirmation of hormone proteoforms by LC-MS/MS islet peptidome

Intact soluble extract from isolated pancreatic islets was fractionated by ultrafiltration (cut-off 10 kDa) and then analyzed by LC-ESI-MS/MS. Ions corresponding to the amylin and C-peptide proteoforms found in MALDI-FTICR were also identified in the islet extracts (**Table 1**), confirming their natural abundance and rejecting the hypothesis for major degradation for these hormones into peptide fragments during MALDI-MS data acquisition.

### Independent intervalidation of hormone peptide proteoforms

A prospective analysis of independent peptidomic data from isolated islets cells was performed using the **MassIVE** database (https://massive.ucsd.edu/ProteoSAFe/massive_search.jsp), which is part of the **Mass Spectrometry ProteinXchange** consortium. This analysis searched for peptide fragments related to the original main pancreatic hormones, restricting the analysis to the sequence of mature peptides. The predictive strategy involved stepwise growing chains from the fixed N-terminus to the C-terminus (*forward*) and from the fixed C-terminus increasing toward the N-terminus (*complementary*). The complete results are sown in **Table S1**.

Eight proteoforms of **amylin** hormone were found, differing in one amino acid for the growing chain, from [1–11]amylin to [1–18]amylin, with the complementary peptide [19–37]amylin also identified (**Table S1**).

The total **glucagon** hormone ([1–29]glucagon) and eight other related shorter proteoforms were identified: [1–26]glucagon, [1–21]glucagon, and [1–13]glucagon to [1–18]glucagon. Glucagon has ^17^Arg, ^18^Arg, and their complementary (*reverse*) fragments were also identified ([18–29]glucagon and [19–29]glucagon, possibly due to the immobilized trypsin used during the proteomic analysis (**Table 2**).

A single *forward* **somatostatin** proteoform was found ([1–12]somatostatin) whereas four *backward-directed* related peptides were identified, two of them after Lysine residues ^14^Lys (^[15–28]^somatostatin) and ^18^ Lys ([19–28]somatostatin).

**Proinsulin-1** and **proinsulin-2** proteoforms were identified in the *forward* form of the entire B-chain plus one or two arginine residues at the C-terminus, indicating a lack of cleavage by proprotein convertase (PC1/3, PC2) and incomplete carboxypeptidase E/H (CPE) removal of the remaining residual c-terminal Arg/Lys. No *reverse* peptides were identified.

Total **C-peptides** from INS-1 and INS-2 were found. No *reverse* peptides derived from **C-peptides** were found. Among the C-peptides, over 13 shorter *forward* proteoforms containing 31 (INS-1) or 29 (INS-2) amino acids were identified. These data suggest the action of a peptidase in a degradation pathway.

The **A-chain** of **INS-1/2** was identified in the complete form and nine *forward* proteoforms. In addition, two *reverse* peptides were found, none after Lys/Arg residues.

The **B-chain** of **INS-1** and **INS-2** was identified in the complete form, and only two shorter peptides (*forward* and *reverse*) were cleaved at ^22^Arg (INS-2), whereas fragments of the remaining chain were not found.

The present findings from other peptidomes in the MASSIVE database provide an intervalidation of our MALDI-MSI analytical approach from histological section, and support the natural abundance of these shorter peptide proteoforms *in vivo*. We interpret these independent findings as evidence for the accumulation of such intermediates in the degradation process of peptide hormones (**Fig. 7**) by islet cell peptidases, which may be due to misfolded or disordered peptide hormones.

## 4. DISCUSSION

Monitoring proteostasis of the endocrine pancreas requires the simultaneous determination of multiple hormones. Immunohistochemical (IHC) techniques, although important tools for quantitative targeted analysis, are limited by several factors, including loss of specificity and sensitivity, tissue fixation/embedding/epitope enhancement procedures ^28,29^, conformational selectivity ^30^, inability to distinguish post-translational modifications, degradation, and proteoforms ^16^. In contrast, molecular histology through spatial proteomic in tissue, as made by MALDI-MSI, enables the identification of known targets, the untargeted discovery and potentially novel biomarkers.

MALDI-MSI has the advantage of spatially resolving the distribution of molecules along a thin histological tissue section providing direct peptidomic information without the need for extraction and proteolysis, albeit with limitations in terms of resolution (instrument-related), abundance, sequencing and low-abundance protein identification. In contrast, it directly detects natural abundance without technical artifacts in peptide recovery during extraction. It has improved positional sensitivity and specificity that limits interference with other adjacent cell types from the same biological sample. For example, islet cells have different hormone production levels that are inversely proportional to the size ^31,32^. Furthermore, peptidomics of individual α-, β-, γ-, and δ-cells, as well as intact islets of Langerhans has shown heterogeneity in cell distribution in mammalian pancreatic islets ^15^. In this study, rat major islet hormone peptides were found either with 2-(4-hydroxyphenylazo)benzoic acid (HABA) for MALDI-MSI in pancreatic thin sections or with 2,5-dihydroxybenzoic acid (DHB) for single-cell profiling MALDI-MS. However, no pancreatic hormone intermediates (precursors/degradation products) were reported, indicating limitations for this purpose ^15^.

Here we report the MALDI-MSI characterization of mice pancreas. Initial experiments were performed with the matrix α-cyano 4-hydroxycinnamic acid (CHCA) (PRENTICE et al., 2019; STEWART et al., 2011). However, few ions were identified using this matrix (**Fig. 2**). Instead, the use of matrix MTB allowed the identification of many endocrine peptides in well-delineated pancreatic islets (**Fig. 4**). In contrast, these peptides were not observed when analyze the whole peptidome from pancreatic extracts (**Fig. 1**). Our MALDI-MSI data also allowed the observation of islet individualities within the same animal, with islets exhibiting individual behavior, with different mass spectra, consistent with different physiological stages ^33,34^.

The optimized method MALDI-MSI allowed direct and selective identification of major hormone proteoforms – shorter peptide fragments - directly from histological sections of the pancreas. We have also identified truncated fragments of hormone peptides, confirmed by LC-ESI-MS peptidomic analysis of islet cell extracts. These fragments were not identified as either whole or isolated cell types in a previous independent study of isolated islets by MALDI-MSI ^15^, indicating the influence of the method on the outcome. Among the amylin fragments we found [19–37]amylin, which was also found in a peptidomic analysis of non-digested islet extracts ^35^. In their study, Boonen and colleagues found no other mature amylin, insulin or glucagon fragments. Instead, they identified many fragments of pro-hormone precursors.

While these fragments could be attributed in part to stress during islet harvest and processing, their detection in our MALDI-MSI data obtained directly from tissue sections suggests that they are most likely endogenous peptides in a truncated form resulting from unexpected cleavage under normal physiological circumstances. The large amount of fragments of C-peptides reported in our and other peptidomic analyzes ^35,36^ is indeed intriguing; they could be involved in several physiological processes, such as peptide degradation pathways, dysfunctional insulin granules, signal transduction, or other endocrine/paracrine functions, as well as protein catabolism and amino acid turnover in nutrient deficiency ^37,38^, which may lead to β-cell failure ^39^.

Eight amylin-related peptides have been identified, ranging from ^[1–11]^amylin to ^[1– 18]^amylin. Mouse amylin has a ^15^Phe that is cleaved by BACE ^40^, originates from membranes of secretory granules and regulates β-cell function and mass ^41,42^, and two arginine residues (^11^Arg and ^18^Arg) that may be associated respectively with [1–11]amylin and [1–18]amylin, when cleaved by trypsin. More than 13 proteoforms of the C-peptides of INS-1 (31 peptides) and INS-2 (29 peptides) were identified. These data suggest the action of multiple peptidases in a degradation pathway, perhaps for *de novo* peptide synthesis.

The exocrine pancreas has several proteases ^43,44^, including endopeptidases (trypsin, chymotrypsin and elastase) and carboxypeptidases such as carboxypeptidase A (selective for aromatic or branched amino acids) and carboxypeptidase B (selective for basic amino acids), which lead to free amino acids and peptide fragments. Other proteases present in the endocrine pancreas are responsible for the conversion of pre-pro-hormones, such as the endoproteases prohormone convertase (PC)1 and PC2, and the zinc-dependent exopeptidase carboxypeptidase H/E (CPH or CPE) ^45^, as well as for degradation processes such as detoxification of misfolded protein or free/excess C-peptide ^46^, including calpain system ^47,48^ and the proteasome pathway ^49^. However, the protease pool of the β-cell (endoplasmic reticulum) consists of endogenous but also exogenous proteases taken up from the exocrine pancreas ^50^.

The secretory granules are acidic compartments ^51^ with a pH of about 5 to 6, consistent with optimal CPH activity ^52^. Zinc-dependent CPH is inhibited by Cu, which is abundant in these granules ^53^. Altered metal homeostasis within β-cells could lead to changes in CPH activity and imbalanced proteostasis that drives lysosomal mediated autophagy, which appears to be necessary for protection against diabetes by clearing misfolded oligomeric β-cell hormones ^54–56^. Thus, the mechanisms for formation of these hormone fragments may involve either biosynthesis or degradation (proteasome/lysosome/autophagy) in the metabolic regulation of nutrients and energy. Other disease or intervention models may be required to further characterize their formation mechanism and putative role.

We compared the control group and the zinc-restricted group using the average spectra of 9 (nine) ROI from 3 (three) animals (3 ROI/animal). The resulting spectra showed a lower signal for the truncated pancreatic peptides in the zinc-restricted animal group compared to the fully mature peptides (insulin, amylin, glucagon). This result could be due to several factors, including modulation of peptide processing by zinc-dependent peptidases, regulation of transcription by zinc, and more. Further transcriptomics and targeted proteomics will contribute to the understanding proteostasis pathways regulated by zinc.

Biomarkers that provide direct evidence of the β-degeneration process have also been monitored. As previously reported, amylin is directly involved in pathophysiology but is not directly related to plasmatic levels. Instead, it accumulated in the pancreas. Although increased production/secretion has been demonstrated by immunohistochemistry in this organ ^6^, such altered levels have not been directly detected in plasma. These data reinforce the importance of histological assessment of the endocrine pancreas, and the need to search for biomarkers that provide direct evidence to identify subclinical stages of diabetes ^5^.

Hyperglucagonemia relative to insulin levels may be a relevant point in the pathophysiology of DM, as there is no pronounced reduction in α-cells in type 2 diabetics ^57^. Instead, mapping of somatostatin (**Fig. 7**) had the lowest resolution and quality among the ions analyzed, indicating its low potential as an indicator of disease progression.

Insulin and amylin are not the only hormones involved in the pathogenesis of DM ^58^. Hyperglycemia in diabetic patients is exacerbated by the inadequately enhanced α-cell function. We have shown that a higher glucagon intensity occurred in the group under dietary zinc restriction. The loss of the balance between insulin and glucagon in T2DM patients may be secondary to the reduction in insulin concentration, suggesting that communication between α- and β-cells may underlie hyperglucagonemia ^59–64^.

The physiological role and regulatory mechanisms of accumulation and action of truncated hormones fragments are unknown. Therefore, further targeted proteomic studies are needed to determine their proteostasis compared with known peptides such as mature insulin and amylin.

## 5. CONCLUSION

This study optimized the matrix-assisted laser desorption/ionization mass spectrometry imaging (MALDI-MSI) technique to analyze mouse pancreatic thin sections. This allowed us to identify the major islet peptide hormones and the accumulated precursors and proteolytic products of these hormones. To validate the identification of these peptide hormones, we performed cross-validation using liquid chromatography-electrospray ionization mass spectrometry (LC-ESI-MS) on islet extracts and an independent peptidome study. Our results showed that dietary zinc restriction reduced the prevalence of truncated hormone forms, suggesting compensatory counter-regulation of peptide degradation, which requires further mechanistic characterization. This novel approach could be further exploited in rodent models to study normal physiology or pathophysiological processes in disease research. With methodological optimization, it may have the potential for pathophysiology research in humans.

## Acknowledgments

We want to thank the staff of the *Centro de Espectrometria de Massa de Biomoléculas* (CEMBIO/UFRJ) and of the *Laboratório Multiusuário de Proteômica* (LMProt/UFMG) for access to their facilities.

## Conflict of Interest

The authors have no financial conflicts of interest with the contents of this article.

## Authorship statement

**TS -** Conceptualization; Data curation; Formal analysis; Investigation; Methodology; Visualization; Roles/Writing - original draft; Writing - review & editing.

**FLB -** Formal analysis; Investigation; Methodology; Visualization; Writing - review & editing.

**DOM –** Methodology; Investigation; Writing - review & editing.

**FSR –** Methodology; Investigation; Writing - review & editing.

**TBM -** Investigation; Writing - review & editing.

**RMB -** Conceptualization; Investigation; Methodology; Supervision; Visualization; Writing - review & editing.

**MTQM -** Conceptualization; Data curation; Formal analysis; Funding acquisition; Investigation; Methodology; Project administration; Supervision; Validation; Visualization; Roles/Writing - original draft; Writing - review & editing.

**LMTRL -** Conceptualization; Data curation; Formal analysis; Funding acquisition; Investigation; Methodology; Project administration; Supervision; Validation; Visualization; Roles/Writing - original draft; Writing - review & editing.

## Funding

This work was supported by the Coordenação de Aperfeiçoamento de Pessoal de Nível Superior (CAPES, Finance Code #001), Conselho Nacional de Desenvolvimento Científico e Tecnológico (CNPq; PQ/311582/2017-6; PQ/313179/2020-4, to LMTRL), Fundação de Amparo à Pesquisa do Estado do Rio de Janeiro Carlos Chagas Filho (FAPERJ; grants E-26/202.998/2017- BOLSA, E-26/200.833/2021-BOLSA, E-26/010.001434/2019-Tematico and E-26/210.195/2020 to LMTRL). The funding agencies had no role in the study design, data collection and analysis, or decision to publish or prepare of the manuscript.

## Data Availability

The data generated during and/or analyzed during the current study are available from the corresponding author on reasonable request.

## Abbreviations

ACN: acetonitrile
DM: Diabetes mellitus
CHCA: α-cyano 4-hydroxycinnamic acid
ESI: Electrospray ionization
FTICR: Fourier Transform Ion Cyclotron Resonance
LC: Liquid Chromatography
MALDI: matrix-assisted laser desorption / ionization MBT, 2-mercaptobenzothiazole
MS: mass spectrometry
MSI: mass spectrometry imaging
TFA: trifluoracetic acid
ToF: time of flight
SPE: solid-phase extraction

---

**Table S1 (spreadsheet). Hormone peptide fragments identified from peptidome database (MassIVE) of isolated islet cells.** Data as obtained using the MassIVE database (https://massive.ucsd.edu/ProteoSAFe/massive_search.jsp), part of the Mass Spectrometry ProteinXchange consortium, using data from ^36^ and others as indicated.

## Notes

### Competing Interest Statement

The authors have declared no competing interest.

